# Functional and Structural Plasticity in Cocaine-Seeking Ensembles of the Nucleus Accumbens Core

**DOI:** 10.1101/2025.10.15.682633

**Authors:** Levi T. Flom, Skylar L. Hodgins, German Gutierrez Erives, Jordan M. Russelavage, Samuel M. Hyken, Zhaojie Zhang, Christopher E. Vaaga, Ana-Clara Bobadilla

## Abstract

Relapse vulnerability in substance use disorder (SUD) is primarily driven by cue-induced activation of neurons within the nucleus accumbens core (NAcore), among other contributing factors. Neuronal ensembles within the NAcore, defined here as selectively co-activated neurons during specific behavioral experiences, are essential during cocaine sensitization and recall. While transient synaptic plasticity (t-SP) has been widely observed in general neuronal populations within the NAcore during reinstatement, its ensemble-specific dynamics remain unclear. Here, we used c-Fos-TRAP2-based tagging to identify cocaine-seeking ensembles in mice following cocaine intravenous self-administration, extinction, and cue-induced reinstatement. Structural spine plasticity was assessed via confocal microscopy, and functional changes were measured using whole-cell electrophysiology across multiple reinstatement time points. Ensemble neurons exhibited enhanced dendritic spine head diameter (d_h_) and AMPA/NMDA (A/N) ratios following cue exposure, consistent with t-SP. Notably, spine classification revealed a reduction in mature spines during reinstatement, suggesting morphological remodeling rather than new spine formation in both ensemble and non-ensemble cells. Non-ensemble neurons exhibited classical functional transient synaptic plasticity, characterized by increased A/N ratios but no significant changes in d_h_. To begin assessing if presynaptic vesicle release impacts t-SP, paired-pulse ratio analysis indicated no differences in population or time point. Importantly, ensemble neurons displayed elevated A/N ratio following cocaine exposure, suggesting prior silent synapse maturation. These findings demonstrate that t-SP is not uniformly distributed across NAcore neurons but differs significantly between ensemble and non-ensemble neurons. By linking ensemble identity to both structural and functional plasticity, this study refines our understanding of cue-induced relapse mechanisms.

**Significance Statement:** Relapse in substance use disorder is strongly driven by cue-induced reactivation of neuronal ensembles in the nucleus accumbens core. While transient synaptic potentiation has been widely described in bulk neuronal populations within the nucleus accumbens core, its ensemble-specific expression has remained unclear. Here, we combined c-Fos-TRAP2 tagging, confocal imaging, and slice electrophysiology to show that transient synaptic potentiation is selectively expressed in behaviorally relevant ensembles. By linking ensemble identity with structural and functional plasticity during cue-induced cocaine seeking, these findings refine current models of relapse and identify ensemble-specific plasticity as a potential target for therapeutic intervention.

## Introduction

Reward-driven behaviors are an essential evolutionary survival strategy, involving neural circuitry in the limbic system (Adinoff, 2004). However, maladaptive alterations within this pathway can lead to substance use disorder (SUD) and drug-seeking behaviors (Koob and Le Moal, 2001; Koob and Volkow, 2010). Within this circuitry, the nucleus accumbens (NAc) serves as a central integration point for reward processing. The NAc receives extensive glutamatergic inputs and is divided into two sub-regions: core and shell (Koob and Volkow, 2010). Importantly, the functional distinctions between the NAc core (NAcore) and shell have prompted researchers to move beyond viewing these regions in isolation, instead considering how populations of medium spiny neurons (MSNs) operate as coordinated neuronal ensembles that drive reward-related behaviors.

Early theories of memory proposed that neurons operate in distributed populations across brain regions to encode and retrieve memories, known as neuronal ensembles (Hebb, 1949). This framework has re-emerged in modern neuroscience (Roy et al., 2022; Williamson et al., 2025), and its application in SUD research has gained momentum (Cruz et al., 2015; Bobadilla et al., 2020; Sortman et al., 2022). One method of identifying neuronal activation within ensembles is the use of immediate early genes, such as c-Fos (Greenberg and Ziff, 1984). Studies have examined the role of c-Fos-expressing neurons in non-contingent cocaine sensitization (Koya et al., 2009; Whitaker et al., 2016) and contingent cocaine self-administration (Chan et al., 2022; Lenoir et al., 2024). Building on this, targeted recombination in active populations (TRAP) (Guenthner et al., 2013), enables endogenous tagging of active cells with the expression of the effector gene tdTomato. TRAP has been used to label the drug-seeking ensemble in the NAcore (Bobadilla et al., 2020; Litif et al., 2023) and other regions (Brown et al., 2023; Wang et al., 2023).

While ensemble tagging reveals which neurons are active, understanding how their synapses adapt requires structural and functional analysis. Many studies have examined dendritic spine plasticity in SUD (Christian et al., 2017; Ku et al., 2024; Mongrédien et al., 2025). One focus is short-term transient synaptic plasticity (t-SP). It is thought to enhance synaptic strength during conditioned cue reinstatement following self-administration of nicotine and cocaine, with increases in AMPA/NMDA (A/N) ratio and spine head diameter (d_h_) leading to glutamate spillover (Gipson et al., 2013a, 2014), which affects both dopaminergic MSN types (Roberts-Wolfe et al., 2018). Additional work on the general population of MSNs in the NAcore shows increases in both A/N ratios and d_h_ following 15-minute cue-induced reinstatement that disappear at 2 hours (Gipson et al., 2013b). Notably, these findings derive primarily from studies conducted in male rats, highlighting the need to consider potential sex differences in t-SP.

A likely mechanism underlying these transient changes is the maturation of silent synapses, spines containing NMDA receptors but lacking AMPA receptors, that form following cocaine use (Whitaker et al., 2016; Wang et al., 2021). Following abstinence, cue exposure can drive the reinsertion of calcium-permeable AMPA receptors (CP-AMPARs) into these spines. However, whether such changes occur specifically within the cocaine-seeking ensemble after self-administration remains unknown.

To address this gap, we utilized the c-Fos-TRAP2 system (DeNardo et al., 2019) to enable long-term tagging of the seeking ensemble in mouse NAcore following cocaine self-administration. We then examined t-SP both structurally, using confocal microscopy, and functionally, using slice electrophysiology, across multiple cue-induced reinstatement time points. We induced expression of an eGFP adenovirus into the NAcore to tag non-ensemble neurons. This allowed the comparison of the structural transient properties of the ensemble with those of non-ensemble neurons in the NAcore. We hypothesized that ensemble neurons would show enlarged spine size and greater A/N ratios compared to their non-ensemble counterparts within the same animals. We show that at 30-minute reinstatement, ensemble neurons have larger d_h_ but similar A/N to non-ensemble neurons. This finding suggests that the previously reported t-SP in non-ensemble neurons at 30 minutes of reinstatement may increase the A/N ratio to ensemble levels during cue-induced reinstatement.

## Methods

### Ai14:c-Fos-TRAP2 Mice

Female (n = 19) and male (n = 31) Ai14:c-Fos-TRAP2 transgenic mice were obtained by crossing male tamoxifen-inducible c-Fos-Cre-recombinase knock-in mice (Fostm2.1(icre/ERT2)Luo/J, Strain# 030323; The Jackson Laboratory) with female Ai14 loxP-flank regulated Cre-reporter knock-in mice (B6;Cg-Gt(ROSA)26Sortm14(CAG-tdTomato)Hze/J; Strain# 007914; The Jackson Laboratory). Mice were individually housed on a 12:12 reverse light schedule during the length of cocaine self-administration conditioning. Sample size and attrition rate were determined from a previously established method (Bobadilla et al., 2020). All procedures were conducted in accordance with the guidelines of the University of Wyoming’s IACUC, Colorado State University’s IACUC, and the NIH’s animal handling procedures.

### Stereotaxic microinjections and catheter implantation

All mice underwent stereotaxic and jugular catheter implantation surgeries at 8-22 weeks of age, performed under isoflurane anesthesia (induction: 3-5% v/v, maintenance: 1-2% v/v). Catheters were inserted 10-12 mm into the jugular vein post-anterior-cervical incision and sutured in place, with the infusion port positioned on the superior cranium using dental cement (CurioDental). Before the installation of the infusion port, 5 µl of non-specific pAAV-hSyn-EGFP (Cat. # 50465-AAV2; Lot # V55182) virus aliquots were diluted from 8.6 x 10^12^ with 15 µl of saline (ICU Medical) for a final concentration of 2.15 x 10^12^ and microinjected 300 nl (AP: +1.5 mm (mice < 25 g) or 1.6 mm (mice > 25 g); ML: ±1.0 mm, DV: 4.4) in the bilateral ventral striatum of c-Fos-TRAP2 mice (n = 34). The injector was then raised by 0.01 mm and left to allow virus dispersal for 8 minutes before being slowly removed from the animal. Mice were then given 4 days to recover before the start of behavioral testing. Recovery was determined by tracking weight gain following surgery, with daily flushes of heparin (1 mg/kg, Fresenius Kabi), intraperitoneal injections of NSAID for analgesia and inflammation (carprofen, 5 mg/kg, MWI), and application of topical triple antibiotics (Medique products). Mice used in functional cohorts received only catheterization and no microinjection (n = 16).

### Cocaine self-administration acquisition, extinction, and reinstatement tests

Both male and female mice were used for self-administration of intravenous cocaine (n = 38) or yoked saline (n = 12) controls (**Figure 1A**). During self-administration acquisition, all mice were on a food-restricted diet (80% of regular food intake = 3.2 g daily) to increase reward-specific operant conditioning responses (Horenstein, 1951; Leslie, 1977; Henton and Fisher, 1981). Female and male 8- to 22-week-old mice (20-30 g) underwent catheter implantation using a previously established method (Bobadilla et al., 2020). All mice underwent self-administration of intravenous cocaine infusions or saline during 2-hour daily sessions for 10 days within standard mouse modular test chambers (Imetronic). Each chamber included two nose-poke (NP) holes associated with either cocaine or saline if yoked saline animal infusions (0.5 mg/kg/inf; NIDA) or inactive NPs. Active NP was paired with an availability light cue (always on until mice interacted with NP; off for ten seconds after NP activation) and an internal NP light cue (on for 3 seconds after NP activation). During sessions, an inactive NP negative control (i.e., with no cues or reward administration) was used to show learning and discrimination. Yoked saline animals were paired with a master cage from one of the cocaine administering cages and would receive an infusion of saline when the master cage received an infusion. For yoked saline animals, the availability light was above the same NP as the master cages’ active NP, but interaction with the active NP by the yoked animal resulted in no change in availability light and did not administer any saline. Following reward-specific cue conditioning, mice underwent a minimum of 10 2-hour extinction sessions in the absence of associated cues, until they were extinguished of rewards via a 70% reduction in active NP association compared to the average of the last three days of cocaine self-administration, for at least the first 30 minutes of an extinction session. At the end of the extinction phase (days 8-10), mice received sham IP saline injections after each session to decrease the response to injections. Following the extinction phase, a 30-minute reinstatement (RST) session with the cocaine-associated light cues and without administering cocaine was conducted. Immediately following the cued reinstatement session, mice received an intraperitoneal injection of 4-hydroxytamoxifen (4-OHT; 50 mg/kg, Sigma)(DeNardo et al., 2019), which activated the Cre-dependent TRAP2 system to label cocaine-seeking ensemble neurons with tdTomato fluorescence. Following initial reinstatement, additional extinction sessions were conducted to allow for fluorescent tdTomato (tdTom) expression into the dendrites and spines, followed by either sacrifice in the homecage (0 timepoint) or a second reinstatement that occurred for 30 minutes or 120 minutes with immediate sacrifice following the reinstatement. All saline control animals were sacrificed from their homecage for sample collection.

**Figure 1.**
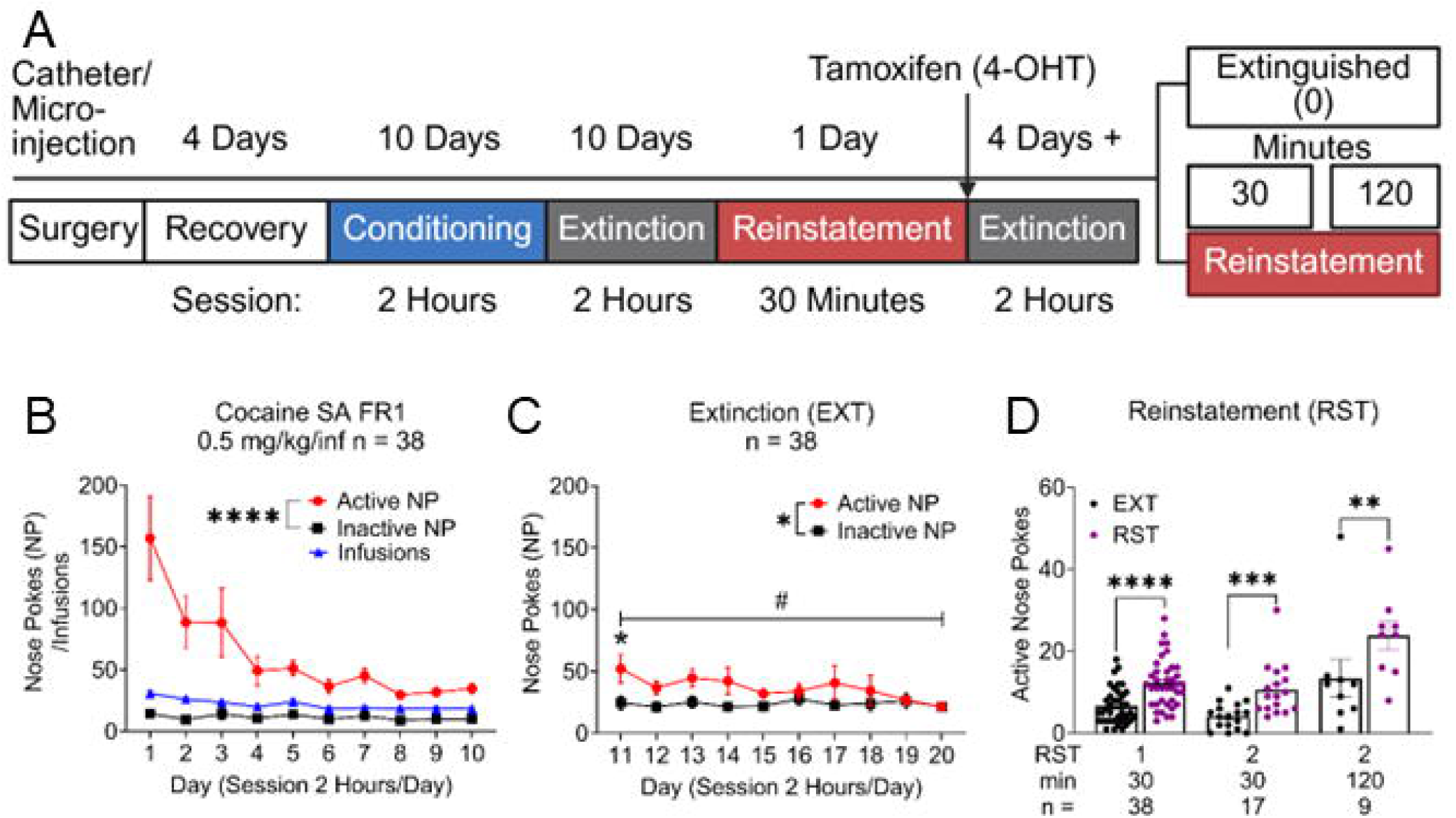
Cocaine self-administration (SA) behavior. (**A**) Cocaine SA timeline. Different colors denote different phases of SA. Animals were injected with 4-hydroxy-tamoxifen (4-OHT) following the first reinstatement (RST). (**B**) c-Fos-TRAP2 mice cocaine SA, **** p < 0.0001 comparing active and inactive nose pokes (NP). (**C**) Extinction (EXT) sessions for mice, * p < 0.05 comparing active and inactive NPs. * p > 0.05 comparing active NP and Inactive NP on day 11, # p > 0.01 comparing active NP on day 11 vs day 20. No difference between active and inactive on day 20, or comparing inactive NPs on day 11 to day 20. (**D**) Active NPs following cue-induced RST for cocaine, **** p < 0.0001 EXT active NP compared to first RST active NP, *** p < 0.001 EXT active NP compared to second RST 30 minutes, ** p < 0.01 EXT active NP compared to second RST 120 minutes.

### Immunohistochemistry

Mice in the structural cohorts from p88-186 were anesthetized with a 2:1 combination of ketamine (MWI) and xylazine (Akorn) and then perfused with 10 mL of 1X phosphate-buffered saline (PBS, Quality Biological) and 20 mL of 3.7% formaldehyde (Fisher Scientific). Brains were post-fixed with 3.7% formaldehyde for at least 24 hours and then transferred to a solution of 20% sucrose, 0.01% sodium azide (Sigma-Aldrich) in 1X PBS. Brains were then sectioned using a cryostat (Leica) at 80 μm to maximize the ability to keep distal dendrites intact. Free-floating sections were then rinsed 3 times in 1X PBS and incubated for 2 hours in a blocking solution of 5% normal goat serum (NGS, Invitrogen), 2.5% bovine serum albumin (BSA, Fisher Scientific), and 0.25% Triton x 100 (Thermofisher) in PBS, then incubated in this blocking solution plus the primary antibody for 12-16 hours to enhance fluorescent signal. Rabbit anti-RFP primary antibody (Rockland Immunochemical, 1:1000 dilution) and chicken anti-GFP primary antibody (Aveslabs, 1:10,000 dilution) were used. The tissue was then rinsed once in 0.25% PBS-Triton, three times in 1X PBS, and incubated for 2 hours in secondary antibodies: Goat anti-rabbit IgG, Alexa Fluor 594 (Thermofisher, 1:1000 dilution), and donkey anti-chicken IgY, Alexa Fluor 488 (Thermofisher, 1:1000 dilution). The tissue was then mounted onto slides using Prolong Gold mounting medium (ThermoFisher).

### Acquisition and quantification of dendritic spines

High-resolution images of 80 µm NAcore slices were acquired using a 63x 1.4 NA, oil immersion objective on a Zeiss LSM 880 (Colorado State University) or 980 (University of Wyoming) confocal microscopes (ZEISS) with voxel size set to 0.164 µm at the XY plane and 0.25 µm intervals along the z-axis with a frame set to 1024x1024 pixels. Medium spiny neurons within the NAcore region, located between Bregma 1.7 mm and 0.62 mm (Paxinos, 2007, 3rd ed.), were identified in Alexa 594 and Alexa 488 laser channels and excited with a 488 nm laser for GFP and a 561 nm laser for tdTomato. Images were converted into Imaris file format using the Imaris file converter. To reduce sample bias, four segments per animal were included for each neuron classification. Using researchers blinded to treatment conditions, dendritic segment images were cropped in Imaris (versions 10.0.0, 10.0.1, Oxford Instruments) starting at least 50 µm from the soma after the first branch point, and segment lengths were 30 - 110 µm for analysis. Following the cropping of the dendritic segments, the images were then cleaned of background using the "surfaces" function in Imaris. Dendritic spines were then identified and quantified using the built-in Imaris machine learning algorithm for filaments programmed with the following parameters: starting point, largest diameter = 1.30 µm, dendrite seed point thinnest diameter = 0.395 µm, diameter of sphere region = 1.30 µm, spine seed point thinnest diameter = 0.217 µm (max length 6.58 µm). Following the creation of the analyzed segments, we then followed up by utilizing the Imaris XT plugin, Classify Spines (Sebastian et al., 2013). We used the following parameters for spine classification, stubby spines: spine length less than 1 um, Mushroom: length of spine less than 3 um and max spine head width greater than mean spine neck width *2, thin spine mean spine head greater than or equal to mean spine neck width, and filipodia/dendrite is all other spines that did not fall into the above category. Note that we only used the classification for stubby, mushroom, and thin spines, but still utilized the filipodia/dendrite-labeled spines for spine density and d_h_ metrics, as they did not fit into any of the previous criteria but were identified as spines by the experimenters.

### Whole cell recordings

Acute brain slices of the NAc for electrophysiology were prepared from p85 to p117 male and female mice. Mice were anesthetized using isoflurane (2-3%) until unresponsive to toe-pinch and then transcardially perfused with ∼10 mL of cold (4°C), oxygenated cutting solution which contained (in mM): 23 NaCl, 3.5 KCl, 1.25 NaHPO_4_, 26 NaHCO_3_, 1 MgCl_2_, 1.5 CaCl_2_, 10 D-glucose (290–310 mosmol, pH 7.3). Coronal slices (250-300 μm), which included the nucleus accumbens, were prepared using Compresstome (VF-510-0Z, Precisionary Instruments). Slices were allowed to recover for at least 30 minutes in warm (37°C) aCSF, which contained (in mM): 123 NaCl, 3.5 KCl, 1.25 NaHPO_4_, 26 NaHCO_3_, 1 MgCl_2_, 1.5 CaCl_2_, 10 D-glucose (290–310 mosmol, pH 7.3).

Whole-cell voltage clamp recordings from both unlabeled and tdTom cells in the NAcore were made using an Olympus BX51WI microscope using CoolLED (550 nm) optical stimulation to identify tdTom cells. To isolate excitatory synaptic transmission, the extracellular solution included 20 μM SR95531 (Tocris Biosciences) to block GABAA receptors. Voltage clamp recordings were made using a Cs-gluconate internal solution, which contained (in mM): 120 CsCH_3_SO3, 3 NaCl, 2 MgCl_2_, 1 EGTA, 10 HEPES, 4 MgATP, 0.3 Tris-GTP, 14 Tris-creatine phosphate, 12 sucrose (288 mosmol, buffered with CsOH to pH 7.32). Borosilicate patch pipettes were pulled to 2–5 MΩ using a Narishige vertical puller. The liquid junction potential (10 mV) was corrected during recording. Temperature was maintained at 35–37°C by a Warner TC-324B controller. Data were digitized at 20 kHz and filtered at 10 kHz. Data was acquired with a Sutter Instruments IPA amplifier/digitizer using the SutterPatch software (IgorPro, Wavemetrics). Before and after data acquisition, access resistance was recorded using a 10-mV hyperpolarizing step, and recordings with changes > 20% across the recording were discarded.

To stimulate cortical inputs to the nucleus accumbens, a bipolar stimulating electrode was placed near the NAcore, generally medial/dorsal to the anterior commissure. To stimulate synaptic transmission, a constant current (0.1 ms, 2-32 mA) was applied to the slice. Synaptic responses were recorded at -70 mV and +40 mV to record both AMPA and NMDA receptor contributions. EPSCs were generated at an inter-stimulus interval of 10 seconds, and ≥10 sweeps at each holding potential were averaged together. Data were analyzed using SutterPatch and IgorPro software. EPSCs at -70 mV and +40 mV were averaged and baseline-subtracted. To estimate the A/N ratio, we measured the peak EPSC amplitude at -70 mV and divided it by the amplitude at +40 mV measured at 40 ms after stimulus onset, to isolate the NMDA receptor component.

### Experimental design and statistical analysis

All statistics were done using GraphPad Prism version 10, and all figures are shown with mean and SEM for error bars. All behavioral data comparisons were analyzed separately at the different phases (acquisition, extinction, and reinstatement), with acquisition and extinction assessed using a two-way Analysis of Variance (ANOVA) with the Geisser-Greenhouse correction. The factors of the analysis were within-subjects for both nose pokes and days. Predetermined paired t-tests were conducted within and between the first and last day of extinction to validate that there was a difference in NP behavior following 10 days of extinction in the chamber. Comparisons of active NPs were conducted between the previous day’s active NP in EXT and the RST active NP with matched time using paired t-tests.

Sex effects were analyzed separately for behavior at the different phases. Acquisition and extinction were assessed using two-way ANOVA with the Geisser-Greenhouse corrections. The factors for the analysis were between-subjects for nose pokes and days. Post-hoc testing was completed using preset comparisons for nose pokes between sexes (i.e., male active NP vs female active NP) and between nose pokes within sexes (male active NP vs male inactive NP).

Yoked saline animals were analyzed with the same paradigm for self-administration and extinction as the cocaine self-administration group to assess for any difference in internal behavior. Reinstatement for the saline animals was evaluated using a paired t-test comparing the first 30 minutes of the last extinction session with the 30 minutes of cued reinstatement. Yoked saline animals were also compared in their active and inactive nose poking to the cocaine self-administration to validate that learning was occurring in the cocaine group using a two-way ANOVA with a within-subjects factor of days, and a between-subjects factor for nose pokes. A separate analysis was done for active and inactive nose pokes between groups. RST data were assessed using a paired t-test design between the previous active NPs during extinction and RST.

Validation of our fluorescent tags was first calculated by taking the average of spine density and d_h_ per channel per segment of neurons tagged with both tdTomato and GFP and then compared using a nested t-test to validate that both fluorescent proteins were suitable for comparison in independent cells. Following this, the average of each segment in both tdTomato-positive (tdTom) and GFP-positive (GFP) cells was merged to assess classical t-SP for spine density and spine diameter (d_h_). A nested ANOVA followed by Tukey’s multiple comparisons test was completed to account for numerous segments from the same animal in the analysis. We then separated the segments by fluorescent tag and reduced our power by calculating the mean of each segment for d_h_, spine density, and the counts of mature and thin spines, and then averaging these values into a single data point per population per animal. This was done to enable the use of two-way ANOVA with a within-subject factor of population and a between-subject factor of time point. Post-hoc tests were conducted with Tukey’s multiple comparison tests. Spine type assays were performed similarly, with two-way ANOVA conducted separately for mature and thin spines. Sex comparisons of structural data were performed using a two-way ANOVA with between-subject factors of sex, population classification and time point as a single variable.

Functional data treated each recorded cell as an independent data point. A/N ratio for cells in animals exposed to cocaine was assessed using an unpaired t-test, with the factor being whether the cell was non-ensemble or ensemble, to investigate if A/N differed following drug exposure between populations. A/N was then assessed with a two-way ANOVA with factors of populations and time point, with Tukey’s multiple comparison tests across timepoints (non-ensemble vs. ensemble), populations across timepoints (saline, 0 minute, 30 minutes), and comparing all time points regardless of population. To assess if presynaptic release influenced cell behavior paired pulse ratio (PPR) was evaluated using the same two-way ANOVA structure.

Regression analysis was conducted for RST2 NPs, examining the size of the d_h_ and A/N ratio to the active NP responding. Two animals were excluded due to catheter failure, one animal was excluded due to a lack of tdTomato expression, and 2 animals were excluded due to loss of cell classification in data from the electrophysiological recordings. Two cells were excluded from A/N recordings (1 ensemble, 1 non-ensemble) due to cell location being in the nucleus accumbens shell, and three were excluded from the non-ensemble group due to outlier testing with ROUT with a Q of 1%. In the PPR data, 24 cells (19 ensemble, 5 non-ensemble) did not respond to the pair pulse stimulus and were excluded from PPR but included in A/N recordings.

## Results

### Cocaine Self-administration, Extinction, and Cue-Induced Reinstatement

To characterize alterations in t-SP both within the cocaine ensemble and non-ensemble neurons, we employed a self-administration (SA) and extinction (EXT) model, using cued reinstatements (RST) at two time points (30 and 120 min), followed by either confocal microscopy imaging or whole-cell electrophysiology. c-Fos-TRAP2 mice underwent daily short-access (2-hour) sessions of intravenous cocaine SA in a Fixed Ratio 1 (FR1) schedule (**Figure 1A**). SA was contingent on cue-paired interaction with a cocaine-reinforced active nose poke (NP). The chamber also included a non-reinforced inactive NP. During all SA sessions, mice interacted more with the active NP than the inactive NP (**Figure 1B**, two-way ANOVA, nose pokes: F (1,37) = 67.20, p < 0.0001, days: F (2.874,106.3) = 6.663, p = 0.0004, interaction F (2.747, 101.6) = 6.141, p = 0.0010). Comparison of male and female revealed an overall effect of time on nose pokes (**Extended Data Figure 1B**, two-way ANOVA, F (9,342) = 8.436, p < 0.0001), no sex differences on active NPs (F (1,38) = 0.7627, p = 0.3880) or interaction (F (9,342) = 0.9514, p = 0.4805). No difference was detected between sex on infusions (**Extended Data Figure 1C**, two-way ANOVA, sex: F (1,38) = 0.6583, p = 0.4222).

Yoked saline control animals followed a similar timeline (**Figure 2A**). During conditioning yoked saline animals received infusions of saline when a paired cocaine animal received an infusion, but no nose poke of the yoked animals lead to an infusion, showed a significant main effect of nose pokes (**Figure 2B**, two-way ANOVA, F (1,11) = 5.885, p = 0.0337), but no difference between days (F (1.809,19.90) = 1.465, p = 0.1716) or interaction (F (1.168,12.85) = 0.4222, p = 0.5582). Comparison of the NPs of the cocaine and yoked saline controls revealed a difference in active NPs between cocaine and yoked saline (**Figure 2C**, two-way ANOVA, days: F (2.7,129.6) = 2.429, p = 0.0747, group: F (1,48) = 12.11, p = 0.0011, interaction: F (2.7, 129.6) = 1.618, p = 0.0011), but no difference in inactive NP (groups: F (1,48) = 0.3446, p = 0.5599, days: F (2.506,120.3) = 2.370, p = 0.0849, interaction: F (2.506,120.3) = 1.040, p = 0.3687).

**Figure 2.**
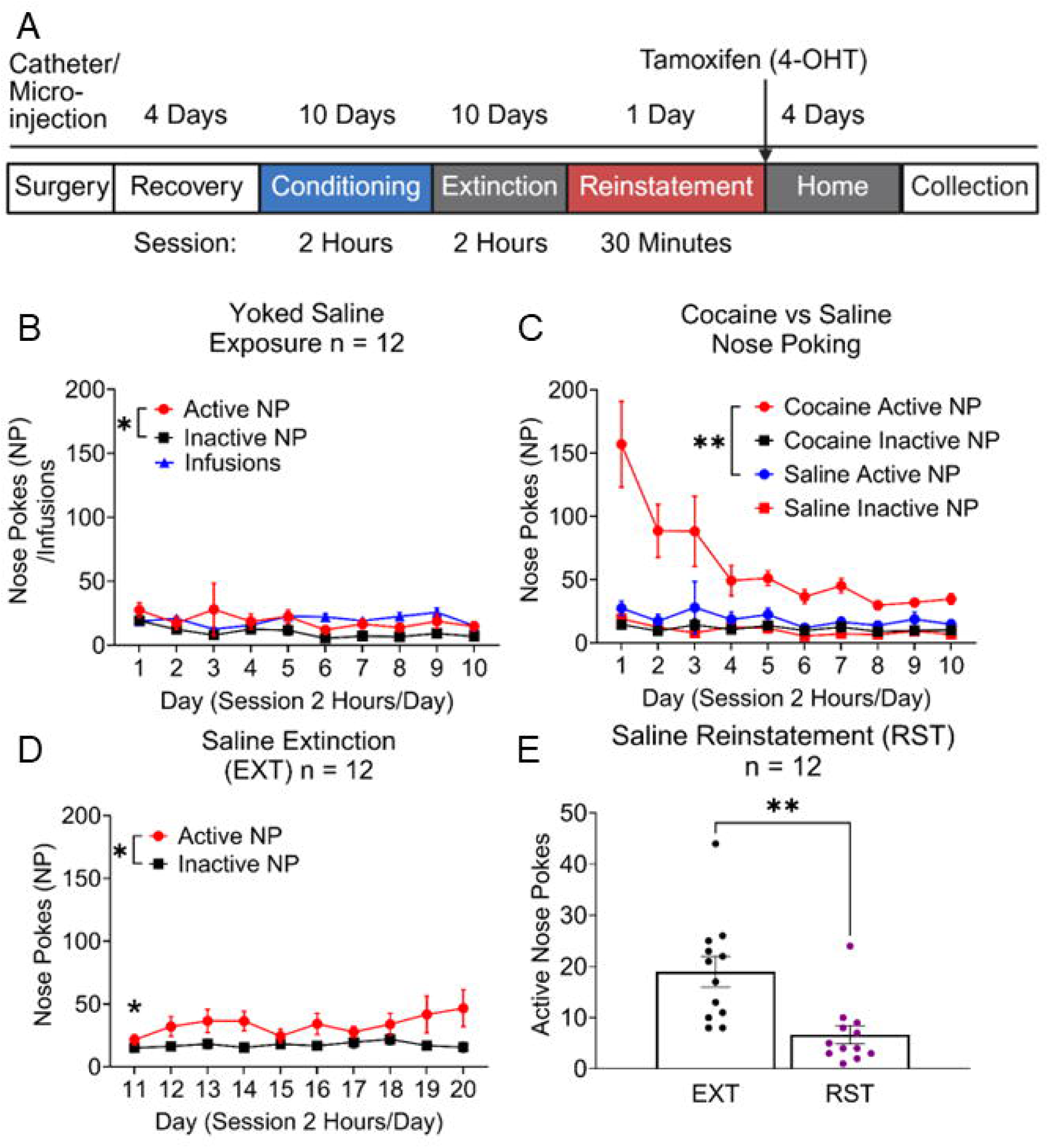
Yoked saline control behavior during self-administration (SA). (**A**) Yoked SA timeline. Different colors denote different phases of SA. Animals were injected with 4-hydroxy-tamoxifen (4-OHT) following the first reinstatement (RST). (**B**) c-Fos-TRAP2 mice behavior during yoked saline conditioning sessions, * p < 0.05 comparing active and inactive nose pokes (NP). (**C**) Comparing nose poking behavior from cocaine exposed animals and yoked controls, ** p < 0.01 saline active NP compared to cocaine active NPs. (**D**) Extinction sessions for mice showed a difference in active and inactive NPs, * p < 0.05, and showed a difference at day 11 that disappeared by day 20. (**E**) Active NPs following cue-induced RST yoked saline, ** p < 0.01 EXT active NP compared to first RST active NP.

Following SA, mice underwent EXT in the original SA chamber without cocaine or related cue exposure. During EXT, mice showed a main effect of NP (F (1.000, 37.00) = 6.702, p = 0.0137), likely due to an extinction burst (planned paired t-test, Day 11 active NP vs inactive NP, t (37) = 2.683, p = 0.0108) but had a non-significant interaction between days and nose pokes (**Figure 1C**, two way ANOVA, F (2.036, 75.35) = 1.622, p = 0.2039). Alternatively, planned comparisons of the NPs between day 11 and 20 revealed a reduction from day 11 to day 20 for active nose pokes (t (37) = 3.072, p = 0.0040), no change in inactive nose pokes (t (37) = 0.5531, p = 0.5835) and showed no difference in nose pokes on day 20 (t (37) = 0.007666, p = 0.9939). This highlights a difference in NPs between active and inactive NPs during the first day of extinction. Still, NP discrimination was lost by day 20, with a reduction in active NPs and no change in inactive nose pokes, leading to no difference on day 20. Male and female comparisons revealed no differences in sex (**Extended Data Figure 1D**, two-way ANOVA, F (1,38) = 0.09777, p = 0.7562), days (F (1.935, 73.52) = 2.531 = 0.0883), or interaction (F (1.935, 73.52) = 1.047, p = 0.3542).

Yoked saline controls animals showed a difference in NPs (**Figure 2D**, two-way ANOVA, F (1,11) = 9.240, p = 0.0113) but no main effect of days (F (2.786,30.65) = 0.6983, p = 0.5503) or interaction (F (2.420,26.62) = 1.362, p = 0.2752). Preplanned comparisons between active and inactive reveal a difference on day 11 (t (11) = 2.234, p = 0.0472), but no difference on day 20 (t (11) = 2.118, p = 0.0578).

To induce cocaine seeking, mice underwent a cued-reinstatement session (**Figure 1D**) where mice were exposed to cocaine specific cues in the absence of cocaine infusions. Mice sought cocaine during the RST1 session as demonstrated with the increase in active NP during RST in comparison to active NP during the same length of time in the EXT session from the day before (**Figure 1D**, paired t-test, t (37) = 5.533, p < 0.0001). Immediately following RST, 4-OHT was administered to tag active c-Fos-dependent neurons in the ensemble underlying cocaine-seeking behavior with tdTomato. Mice were then given 5 days of additional EXT (data not shown) to allow for expression of tdTomato to fill the dendrites and spines. They were then either sacrificed in the home cage (0 minutes) or sacrificed immediately following a second cued reinstatement at 30 minutes (**Figure 1D**, paired t-test, t (16) = 4.324, p = 0.0005) or 120 minutes (**Figure 1D**, paired t-test, t (8) =3.698, p = 0.0181). Males showed significant reinstatements across all time points (**Extended Data Figure 1E**, EXT1 vs RST1: paired t-test, t (23) = 5.152, p < 0.0001, EXT2 30 minute vs RST2 30 minutes t (11) = 6.124, p < 0.0001, EXT2 120 vs RST2 120 t (5) = 3.538, p = 0.0166) while females showed a significant first reinstatement (EXT1 vs RST1, t (13) = 2.528, p = 0.0252) and a non-significant trend toward an increase in second reinstatements (EXT2 30 minutes vs RST2 30 minutes: t (4) = 2.116, p = 0.1017, EXT2 120 minutes vs RST2 120 minutes: t (2) = 1.352, p = 0.3091). Comparing the RSTs between sexes revealed no differences at any time point (unpaired t-test RST1: t (36) = 0.3065, p = 0.7618, RST2 30 minutes: t (15) = 2.127, p = 0.0504, RST2 120 minutes: t (7) = 0.4682, p = 0.4682).

Saline animals underwent cued reinstatement for 30 minutes, revealing a significant reduction in nose pokes during reinstatement (**Figure 2E**, paired t-test, t (11) = 3.963, p = 0.0022), confirming that no association formed with the active NP and saline infusions.

Following euthanasia, mice underwent either transcardiac perfusion for structural analysis or slice preparation for functional analysis. The presence of tdTom fluorescence identified neurons that were part of the ensemble. The non-ensemble neurons were characterized by the presence of GFP in the soma and dendrites of MSNs in structure cohorts or the absence of tdTom in functional cohorts. Following the identification of tdTom for ensemble classification and verification of location in relation to the anterior commissure, we proceeded with structural and functional analysis.

### TdTomato and GFP tagging are identical for spine density and spine head diameter metrics

To ensure that the fluorescent tags of both the tdTom ensemble and GFP non-ensemble neurons were comparable, we compared structural measurements in double-labeled tdTom and GFP cells. We confirmed that both tags fill the dendritic spines equally (**Figure 3A**). Comparing the two channels revealed no differences in spine density (**Figure 3B**, nested t-test, t (18) = 0.7007, p = 0.4924) or d_h_ (**Figure 3C**, nested t-test, t (18) = 0.8343, p = 0.4150). We also ensured that microinjection targets were positioned within the NAcore (**Extended Data Figure 2**).

**Figure 3.**
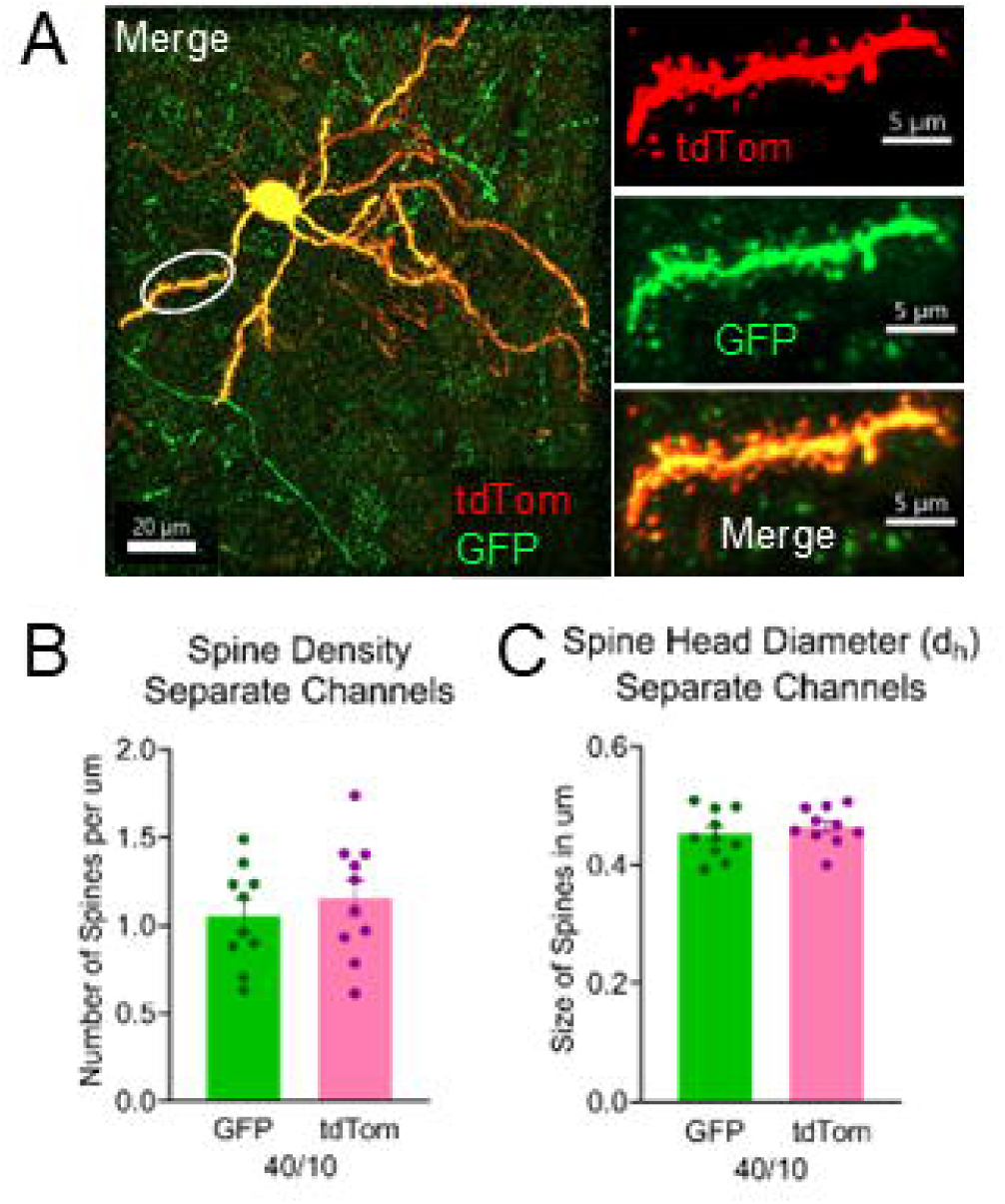
Validation of fluorescence tags being comparable. (**A**) Representative image of a neuron that expresses tdTom (red) and GFP (green) (left). Split of channels (right) showing spines in isolated channel, tdTomato (top), and GFP (middle), merged image of a dendritic segment showing both channels (bottom). (**B**) Spine density on overlapping dendritic segments shows no difference based on channel. (**C**) Spine head diameter (d_h_) on overlapping dendritic segments shows no difference based on channel. Numbers below denote segments over the number of animals.

### Neurons show t-SP following 30 minutes of secondary reinstatement, which is enhanced in ensemble neurons

Following validation and quantification of segments using Imaris (**Figure 4A**), we assessed t-SP in the pooled population (ensemble tdTom and non-ensemble GFP neurons) to test if we could replicate previous results measuring t-SP in all cells within the NAc (Gipson et al., 2014). We observed an increase in spine density following exposure to cocaine (**Figure 4B**, nested one-way ANOVA, F (3,30) = 9.564, p = 0.0001) with Tukey’s post-hoc testing revealing a significant increase at 0 minute (p = 0.0010), 30 minutes (p = 0.0356), but not at 120 minutes (p > 0.999) compared to saline. We also observed differences between the 120-minute animals and the 0-minute animals (p = 0.005) or the 30-minute group (p = 0.0232). We also assessed d_h_ and found increases following cued reinstatement for cocaine (**Figure 4C**, nested one-way ANOVA, F (3,30) = 4.008, p = 0.0164). Post-hoc analysis revealed an elevation in d_h_ from the saline animals following both 30-minute (p = 0.0251) and 120-minute ( p = 0.0304) RST2 but no elevation from cocaine extinguished 0-minute animals (p = 0.5659), as well as no differences between the cocaine exposed animals across time points (0 minute vs 30 minutes p = 0.4088, 0 minute vs 120 minutes p = 0.4210, 30 minutes vs 120 minutes, p > 0.9999). We then separated the segments between non-ensemble and ensemble cells, as well as the time point of collection (0 minutes, 30 minutes, or 120 minutes). We found no significant main effect in ensemble vs non-ensemble neurons for spine density (**Figure 4D**, two-way ANOVA, F (1,30) = 1.217, p = 0.2787). However, we noted significant differences across time points in spine density (F (3,30) = 9.564, p = 0.0001) and a significant interaction (F (3,30) = 3.946, p = 0.0174). Tukey’s multiple comparisons reveal that ensemble 0-minute neurons have a greater spine density compared to saline ensemble controls (p < 0.0001), ensemble 30-minute RST2 (p = 0.0152), and 120-minute RST2 ensemble neurons (p < 0.0001). Ensemble 30-minute neurons were shown to be significantly different from saline ensemble neurons (p = 0.0460) and from their non-ensemble counterparts (p = 0.0236). No differences were observed in the non-ensemble population for spine density. Comparing males and females revealed no changes in spine density between sexes (**Extended Data Figure 3A**, two-way ANOVA, F (1,52) = 0.04343, p = 0.8357).

**Figure 4.**
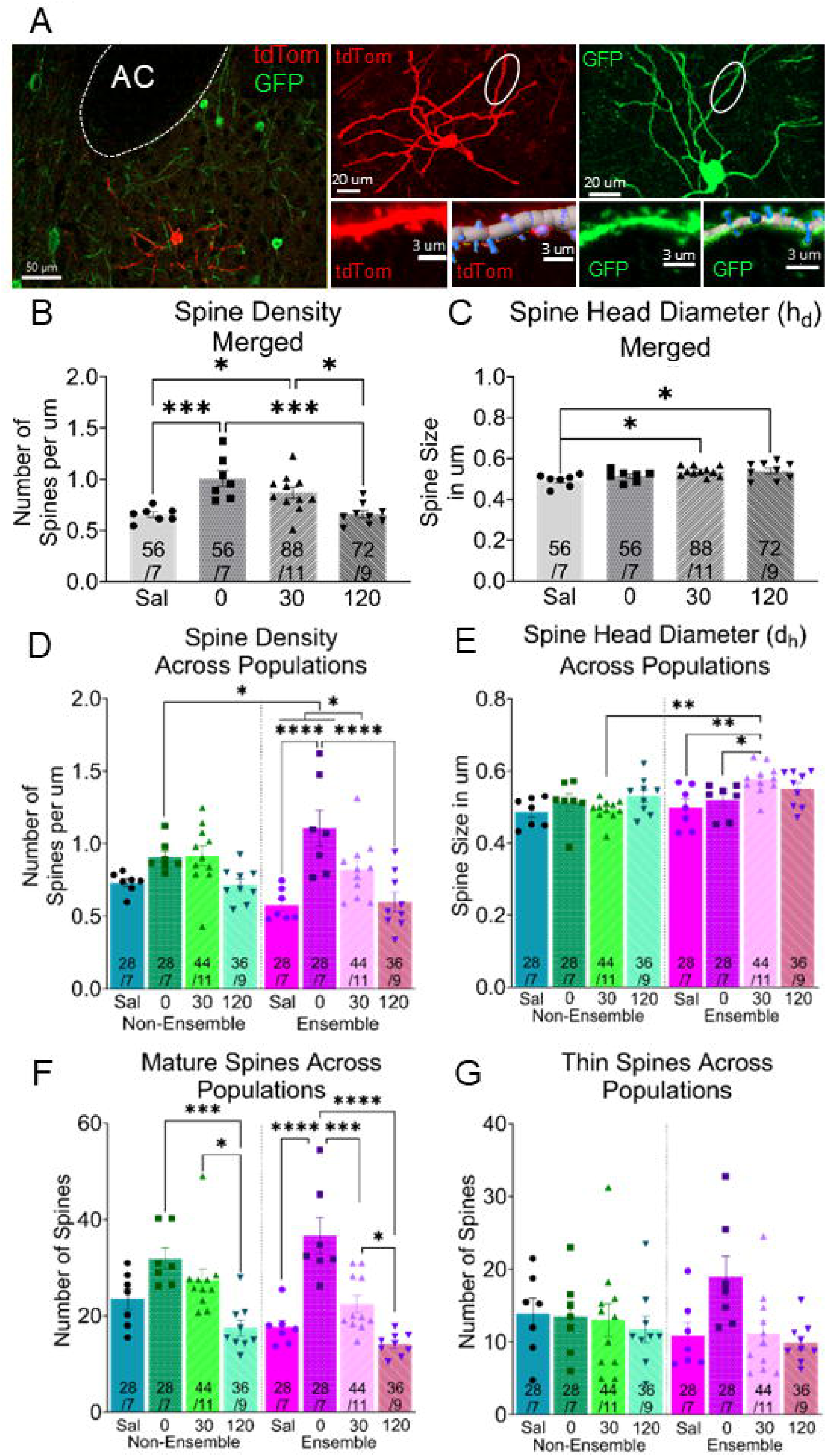
Structural characteristics of non-ensemble and ensemble neurons. (**A**) Left: representative images of nucleus accumbens core with non-ensemble GFP (green) and ensemble tdTom (red) neurons. Center: tdTom neuron with measured segment indicated (top), with segment cropped (left) and segment after analysis (right). Right: GFP neuron with measured segment indicated(top), GFP segment cropped (left), and after analysis(right). AC, anterior commissure. (**B**) All ensemble and non-ensemble dendritic segments together (merged) display changes in spine density. *** p < 0.001, yoked saline (Sal) and 0-minute reinstatement (RST2) segments, * p < 0.05 saline and 30-minute RST2 segments. * p < 0.05 30-Minute and 120-minute RST2 segments. *** p < 0.001 0-minute and 120-minute RST2 segments. (**C**) Merged dendritic segments exhibit changes in spine head diameter (d_h_). * p < 0.05 Sal compared to 30-minute and 120-minute RST2. (**D**) Spine density across behavioral groups, Ensemble 0-minute RST2 compared to yoked saline, 120 minutes, **** p < 0.0001. * p < 0.05, Ensemble 30 minutes compared to Sal and 0-minute ensemble RST2 segments. * p < 0.05 0-minute ensemble and 0-minute non-ensemble segments. (**E**) d_h_ Across behavioral groups. 30-minute in the ensemble compared to ensemble yoked saline and 30-minute non-ensemble RST2 segments, ** p < 0.01, 0-minute ensemble and 30-minute ensemble RST2 segments, p < 0.05. (**F**) Mature spines across behavioral groups. Non-ensemble 0-minute compared to 120-minute RST, *** p < 0.001. Non-ensemble 30-minute compared to 120-minute RST, * p < 0.05. Ensemble 0 minute compared to Sal, 120-minute RST, **** p < 0.0001, Ensemble 0 minute compared to 30 minute, *** p < 0.001, Ensemble 30-minute RST2 compared to 120-minute RST2 * p < 0.05. (**G**) Thin spines across behavioral groups. Numbers represent segments over the number of animals.

Examining changes in d_h_ revealed differences between ensemble and non-ensemble neurons (**Figure 4E**, two-way ANOVA, F (1,30) = 7.316, p = 0.0112), time points (F (3,30) = 4.006, p = 0.0164), and interaction (F (3,30) = 3.019, p = 0.0452). Multiple comparisons revealed that ensemble spines at the 30-minute timepoint are significantly larger than ensemble yoked saline (p = 0.0024), 0-minute ensemble (p = 0.0222), and 30-minute non-ensemble segments (p = 0.0002). This change in d_h_ was not seen in non-ensemble comparisons. Comparisons of males and females revealed no difference in d_h_ based on sex (**Extended Data Figure 3B**, two-way ANOVA, F (1,52) = 2.251, p = 0.1396).

### Spine classification reveals similar trends to mature spine destabilization across ensemble and non-ensemble neurons

Using the Imaris XT spine classification plugin enabled us to categorize spines into two classifications: “mature” (characterized by stubby and mushroom spines) and “immature” (characterized by thin spines). Analysis of mature spines reveals a main effect between timepoints (**Figure 4F**, two-way ANOVA, F (3,30) = 20.78, p < 0.0001) but no main effect for ensemble vs non-ensemble (F (1,30) 2.263, p = 0.1430) or interaction (F (3,30) = 2.151, p = 0.1147). Non-ensemble neurons showed a reduction in mature spines from 0 minutes to 120 minutes RST2 (p = 0.0001) and from 30 minutes RST2 to 120 minutes RST2 (p = 0.0138). Ensemble neurons show an increase in mature spines at 0 minutes compared to yoked saline ensemble neurons (p < 0.0001). The number of mature spines decreases from 0 minutes following 30-minute RST2 (p = 0.0001) and 120-minute RST2 (p < 0.0001), as well as from 30 minutes ensemble to 120 minutes non-ensemble (p = 0.0270). Thin spines, however, do not display these traits (**Figure 4G**, two-way ANOVA, ensemble vs non-ensemble: F (1,30) = 0.2964, p = 0.05902, timepoint: F (3,30) = 1.778, p = 0.1726, interaction: F (3,30) = 2.479, p = 0.0803). Comparing sexes revealed no differences in spine classification (**Extended Data Figure 3C-D**, two-way ANOVA, F (1,104) = 2.865, p = 0.0935).

We aimed to determine whether the levels of seeking during RST2 correlated with the spine size of ensemble and non-ensemble segments at 30 minutes and 120 minutes. We found no significant correlation for non-ensemble or ensemble d_h_ at 30 minutes (**Extended Data Figure 4A**, simple linear regression, non-ensemble, R^2^ = 0.008296, p = 0.5565, ensemble R^2^ = 0.02914, p = 0.2679) or 120 minutes (**Extended Data Figure 4B**, simple linear regression, non-ensemble, R2 < 0.0001, p = 0.9761, ensemble R^2^ = 0.08297, p = 0.6560).

### Ensemble-specific functional glutamatergic plasticity

To better understand potential functional changes, we employed whole-cell patch clamp electrophysiology in the NAcore (**Figure 5A**) from ensemble and non-ensemble neurons. More specifically, we recorded the changes in the AMPA and NMDA receptor ratio (A/N) from saline control, 0-minute, and 30-minute RST2 animals (**Figure 5B**). Merging ensemble and non-ensemble cells reveals a main effect of time points on A/N ratio between time points (**Figure 5C**, one-way ANOVA, F (2,85) = 8.225, p = 0.0005). Multiple comparisons reveal that 30-minute RST2 shows elevated A/N compared to saline (p = 0.0011) and 0-minute animals (p = 0.0063). We then separated ensemble and non-ensemble neurons in cocaine exposed animals. We found that following cocaine exposure, ensemble neurons exhibit a greater A/N ratio compared to non-ensemble neurons, regardless of reinstatement status. Indicating altered excitatory balance within ensembles following cocaine exposure (**Figure 5D**, unpaired t-test, t (56) = 2.478, p = 0.0163). Lastly, we separated the cells into ensemble vs non-ensemble and time point groups. We found a main effect of time point (**Figure 5E**, two-way ANOVA, F (2,82) = 8.540, p = 0.0004) but no main effect for ensemble vs non-ensemble (F (1,82) = 2.612, p = 0.1099) or interaction (F (2, 82) = 1.069, p = 0.3481). Multiple comparisons reveal that in non-ensemble cells, A/N increases occur from 0 minutes to 30 minutes (p = 0.0330) but not from saline to 30 minutes (p = 0.1323). Qualitatively, ensemble neurons appeared to show a smoother increase in A/N ratio across the three timepoints, whereas the non-ensemble neurons showed a more abrupt increase in A/N at the 30-minute timepoint. In ensemble neurons, we see an increase in A/N from saline to 30-minute RST2 (p = 0.0021). We also noted a trending rise in difference of A/N between non-ensemble and ensemble neurons, specifically at the 0 timepoint (p = 0.0616) with ensemble neurons having increased A/N. When comparing males and females, there was no difference in ensemble (**Extended Data Figure 4C**, two-way ANOVA, F (1,41) = 0.4800, p = 0.4923) or non-ensemble cells (F (1,38) = 1.007, p = 0.3220).We then wanted to confirm whether the behavior on RST2 correlated with differences in A/N ratios (**Extended Data Figure 4D**). Comparing A/N to 30-minute RST2 animals revealed no correlation in non-ensemble (simple linear regression, R^2^ = 0.03698, p = 0.4923) and ensemble (R^2^ = 0.001718, p = 0.8703) neurons and A/N.

**Figure 5.**
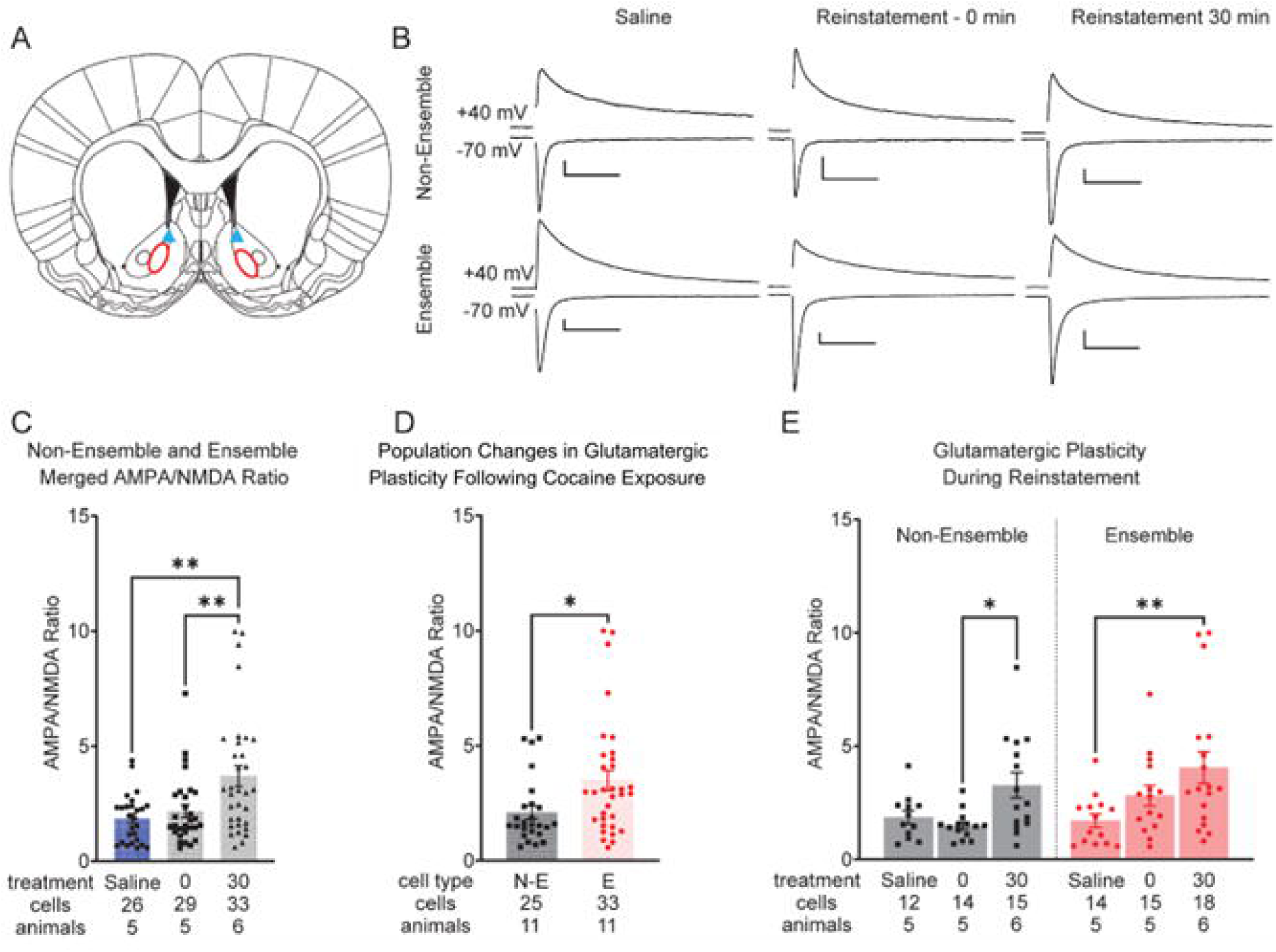
Functional plasticity alters following cocaine exposure. (**A**) Schematic of stimulating electrode (blue triangle) and patching area (red circle) for the nucleus accumbens core. (**B**) Representative traces of all behavioral groups, AMPA (top) and NMDA (Bottom). Scalebars 100 pA / 20 ms (**C**) AMPA to NMDA ratios (A/N) were significantly elevated from cocaine self-administration compared to saline control animals, p < 0.01. (**D**) Ensemble neurons (E) have elevated A/N compared to non-ensemble neurons (N-E) after cocaine exposure, * p < 0.05. (**E**) Cue-induced reinstatement elevates A/N ratios in both non-ensemble and ensemble neurons within cell type but not between groups, non-ensemble 0-minute compared to non-ensemble 30-minute RST, * p < 0.05, ensemble yoked Sal compared to ensemble 30-minute RST2, ** p < 0.01.

To begin to assess whether synaptic release might differ between ensemble and non-ensemble neurons in the NAcore during cue-induced reinstatement, we utilized PPR to examine potential differences across time and cell classification. While PPR is an indirect correlative measure rather than a direct index of release probability, it provides a useful comparative metric under these conditions. We found that there was no difference between ensemble and non-ensemble neurons (**Extended Data Figure 5**, two-way ANOVA, F (1,63) = 0.07784, p = 0.7812), timepoint (F (2,63) = 0.06269, p = 0.9393), or an interaction between the two (F (1,63) = 0.07784, p = 0.4836).

## Discussion

Using confocal microscopy and whole-cell electrophysiology, we expanded the understanding of t-SP in the NAcore and characterized the plasticity of ensemble and non-ensemble neurons during cue-induced reinstatement. Following successful behavioral responding, we confirmed the classic expression of t-SP in mice, and that ensemble neurons show greater structural changes than non-ensemble counterparts at peak t-SP (30 min). Next, we classified the spine data into types based on shape classification. We found that mature spines were less frequent over the course of t-SP regardless of whether the cell was ensemble or non-ensemble. Finally, we observed that neurons exhibited increases in the A/N ratio consistent with t-SP and that ensemble neurons showed a greater A/N ratio compared to non-ensemble neurons following cocaine exposure regardless of the timepoint of reinstatement. These differences are more consistent with postsynaptic alterations, as we did not detect accompanying changes in PPR, although PPR provides only an indirect measure of presynaptic vesicle release. Taken together, these data reveal the importance of both non-ensemble and ensemble-specific t-SP in cocaine seeking, in accordance with the hypothesis that ensemble neurons would show greater attributes of t-SP, including d_h_ and A/N ratio.

### Ensemble-specific synaptic remodeling links silent synapse maturation to transient synaptic plasticity during cue-induced cocaine reinstatement

A large body of research has identified neuronal ensembles, sparsely distributed populations of neurons activated during specific behavioral experiences, as critical mediators of drug-seeking and relapse (Koya et al., 2009; Bossert et al., 2011). Using activity-dependent tagging tools, such as Fos-GFP rats, prior work has demonstrated that ensemble neurons in the NAc are essential for reinstatement behavior and exhibit unique molecular adaptations following drug exposure (Koya et al., 2012). One adaptation is the elevated formation of silent synapses, glutamatergic synapses that contain NMDA receptors but lack functional AMPA receptors (Huang et al., 2009). Silent synapses are believed to be a key form of metaplasticity, priming ensemble neurons for later strengthening during re-exposure to drug-associated cues (Koya et al., 2012; Whitaker et al., 2016). Despite these insights, the temporal evolution of synapse-associated plasticity in ensembles, particularly during withdrawal and reinstatement, remains incompletely understood, as ensemble studies have primarily employed immediate tagging and retrieval for analysis with Fos-GFP rats that allow for peak ensemble tag at 90 minutes post behavior but inability to study how these ensemble neurons change in AMPA or NMDA over extended periods (Whitaker and Hope, 2018). We hypothesize that the changes observed in the A/N ratio between ensemble and non-ensemble cells are partly due to the maturation of silent synapses.

Independent of ensemble identity, several studies have characterized the time-dependent maturation of silent synapses during cocaine withdrawal (Lee et al., 2013; Wright et al., 2020). Early withdrawal (e.g., 24–48 hours) is associated with an increase in silent synapses in the NAc, particularly within amygdala or cortical projections (Huang et al., 2009; Lee et al., 2013). Over prolonged withdrawal periods, these silent synapses typically undergo maturation, incorporating CP-AMPARs and thus transforming into active, strengthened excitatory inputs (Lee et al., 2013; Wang et al., 2021). This maturation is believed to contribute to the incubation of craving, a phenomenon where cue-induced drug seeking increases over time (Wolf, 2016). This incubation of craving has been shown to persist behaviorally for up to 90 days (Grimm et al., 2003) and has been studied structurally for up to 28 days of withdrawal (Graziane et al., 2016). However, these studies do not resolve whether silent synapse formation and maturation occur specifically within ensemble neurons, those that are reactivated during cue-induced relapse, or whether these processes occur globally across the NAc. The present study falls into intermediate withdrawal, being 14+ days removed from the last administration of cocaine.

Parallel work on synaptic changes has focused on t-SP as a circuit-level mechanism of cue-induced relapse (Shen et al., 2011; Gipson et al., 2013a). Upon exposure to drug-paired cues, NAcore neurons exhibit a rapid, transient increase in d_h_, A/N ratio, and glutamate spillover, peaking around 15–30 minutes after cue exposure and resolving within two hours. This form of plasticity is temporally linked to drug-seeking behavior and has been widely replicated across cocaine and nicotine (Gipson et al., 2013a, 2013b; Garcia-Keller et al., 2020; Roberts-Wolfe et al., 2020). However, these findings reflect changes in the bulk population of NAcore neurons and do not account for ensemble recruitment. Whether t-SP occurs preferentially in behaviorally activated ensembles or reflects a population-wide adaptation has remained an open question that we addressed in this project.

The present study helps bridge this gap by showing that t-SP occurs preferentially within ensemble neurons in the NAcore during cue-induced reinstatement. Using TRAP2-based tagging and confocal imaging, we found that ensemble neurons, but not non-ensemble counterparts, exhibit increased d_h_ following 30 minutes of reinstatement. Importantly, we utilized both male and female mice, which may account for our failure to replicate prior reports that t-SP shows d_h_ enlargement in general population. These changes are consistent with the structural hallmarks of t-SP and suggest that previously reported population-level findings may have been driven primarily by adaptations in ensemble neurons. Furthermore, spine classification analysis revealed a reduction in mature spines following reinstatement, without a corresponding increase in thin spines, suggesting morphological remodeling of large spines rather than the generation of new immature ones during reinstatement that occurs in both ensemble and non-ensemble neurons. This pattern is consistent with the hypothesis that silent synapses formed earlier in drug experience may persist or re-emerge in ensemble neurons and undergo dynamic maturation during relapse (Lee et al., 2013; Wang et al., 2021).

Functionally, we showed that ensemble neurons have a greater ratio of A/N following cocaine exposure and withdrawal, likely due to silent synapse maturation after initial ensemble formation (Figure 5D) (Wright and Dong, 2021). This difference in AMPA-mediated signaling revealed no transient increase post-cued reinstatement compared to 0-minute extinguished animals for ensemble cells, but an increase compared to drug naïve ensemble cells. Non-ensemble neurons increase AMPA signaling following 30 minutes of cue-induced reinstatement, replicating previous literature showing an increase in AMPA-mediated signaling in cocaine rats (Gipson et al., 2013a; Roberts-Wolfe et al., 2020). Importantly, these changes were not accompanied by alterations in PPR, indicating that postsynaptic mechanisms, rather than changes in presynaptic release probability, underlie the observed plasticity.

### Study Limitations and Sex Differences

While present findings indicate that ensemble neurons are the primary locus of reinstatement-related plasticity, we also observed that non-ensemble neurons exhibit delayed or muted changes, consistent with a background level of plasticity or spillover from the broader circuit (Gipson et al., 2014). These findings support a model in which t-SP is not a uniform adaptation across NAcore neurons but is instead compartmentalized within a functionally defined subset of neurons that were reactivated by drug-associated cues. Notably, our study did not directly track the formation or fate of silent synapses in ensembles across withdrawal periods. However, the structural data, combined with existing literature, support the possibility that ensembles may retain a capacity for rapid, cue-triggered spine remodeling, potentially mediated by the maturation of previously silent synapses. We also examined sex as a biological variable and observed no significant differences in structural or functional measures across male and female mice. However, the limited female sample size due to breeding constraints warrants caution in interpretation. Given the well-documented sex differences in drug-seeking behavior and relapse susceptibility (Lynch and Carroll, 2000; Hilderbrand and Lasek, 2014). Future work should be designed with sufficient power to detect potential sex-specific effects in ensemble plasticity.

## Conclusions

Together, these findings offer new insight into the cellular specificity of synaptic plasticity during cocaine relapse. By demonstrating that ensemble neurons exhibit classical t-SP features during cue-induced reinstatement, the present study integrates previously distinct models of ensemble function and synaptic plasticity. This ensemble-focused lens refines general understanding of relapse mechanisms, suggesting that therapeutic strategies targeting ensemble-specific plasticity rather than global population modulation may yield greater specificity and efficacy in preventing relapse.

## Data Availability Statement

The raw and analyzed data that support the findings of this study are available from the corresponding author upon reasonable request.

## Acknowledgments

The authors thank the NIDA Drug Supply Program, administered by the Division of Therapeutics and Medical Consequences, for providing cocaine. They also thank the University of Wyoming’s Integrated Microscopy Core (IMC) headed by Zhaojie Zhang for their technical assistance and all members of the Bobadilla Lab for their help and support.

---

**Extended Data Figure 1.**
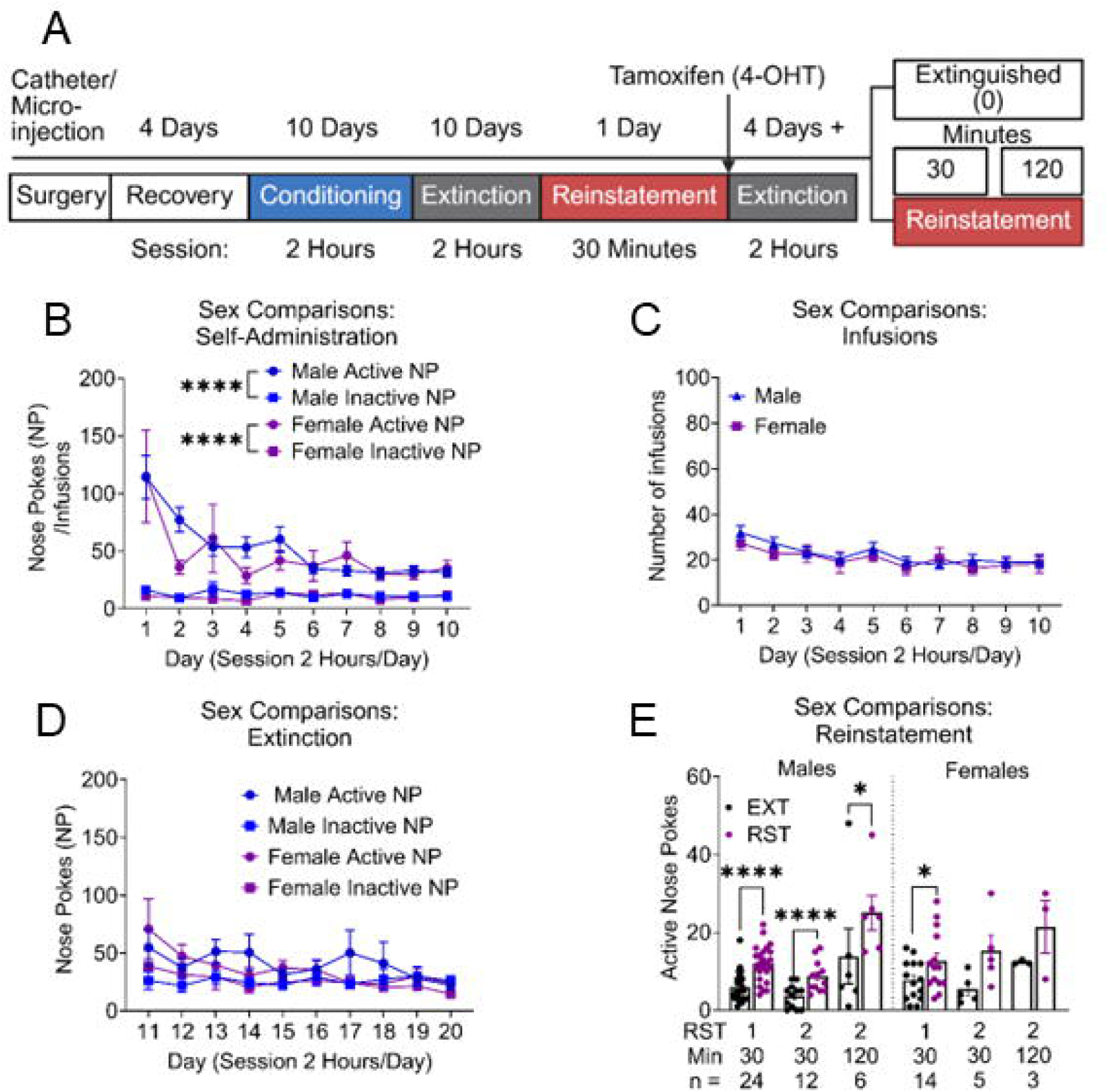
Male and female c-Fos-TRAP2 mice during cocaine self. administration (SA) behavior. (**A**) Cocaine SA timeline for males and females. Different colors denote different phases of SI\. Animals were injected with 4. hydroxy-tamoxifen(4-0HT) following the first reinstatement (RST). (**B**) c-Fos­ TRAP2 mice cocaine SA, no difference between sex on active and inactive nose pokes(NP). **** p < 0.0001 Active vs Inactive NP within sexes. (**C**) Infusions between sexes display no difference in intake. (**D**) Extinction sessions showed no difference in sex on NPs. (**E**) Active NPs following cue-induced reinstatement (RST) for cocaine was elevated only with male mice with no difference between sex on RST level, **** p < 0.0001 male EXT active NP compared to first RST active NP, ••• p < 0.001, male EXT active NP compared to second RST 30 minutes, • p < 0.05, male EXT active NP compared to second RST 120 minutes.

**Extended Data Figure 2.**
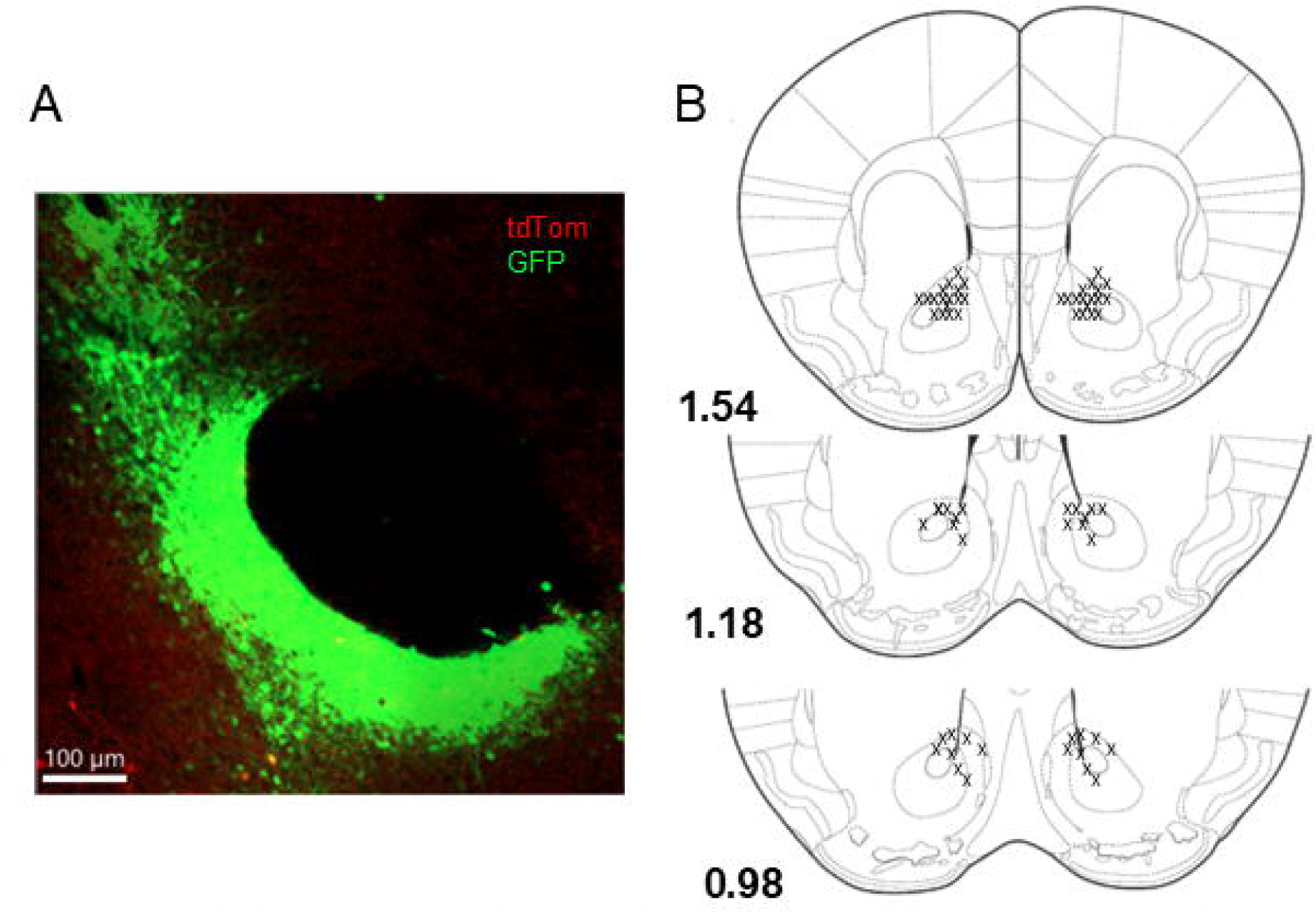
Virus tract locations in the nucleus accumbens core. (**A**) Representative 10x image of GFP virus trad and injection site (**B**) atlas images of virus injection sites. X denotes virus injection sites from 34 animals, modified from Paxinos and Franklin’s mouse brain in stereotaxic coordinates, 5^th^ edition.

**Extended Data Figure 3.**
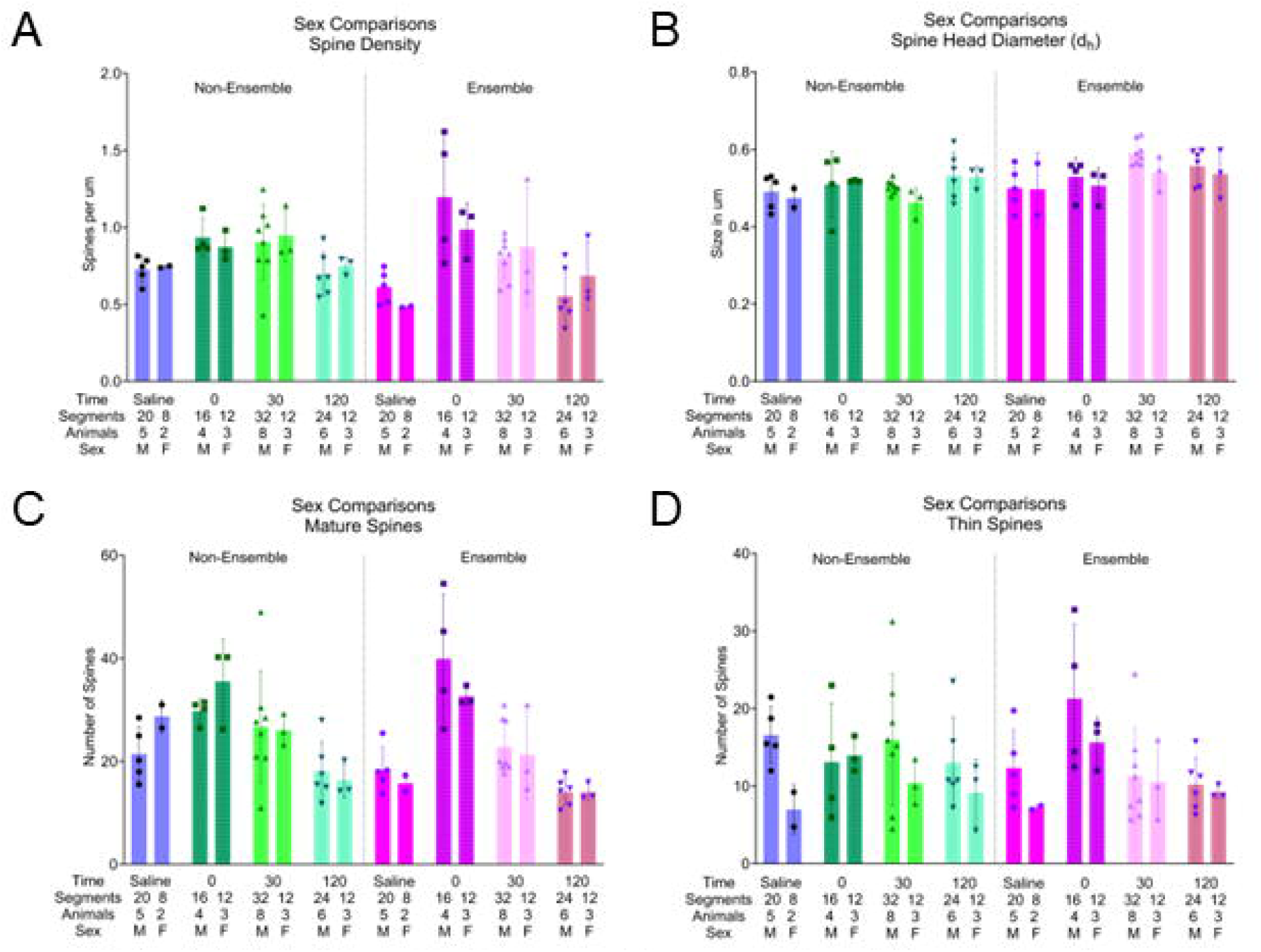
Sex comparisons reveal no difference in spine morphology ( **A**) Spine density does not differ across sexes. (**B**) Spine head diameter does not vary across sexes. (**C**) The number of mature spines does not vary across sexes. (**D**) The number of thin spines does not vary across sexes.

**Extended Data Figure 4.**
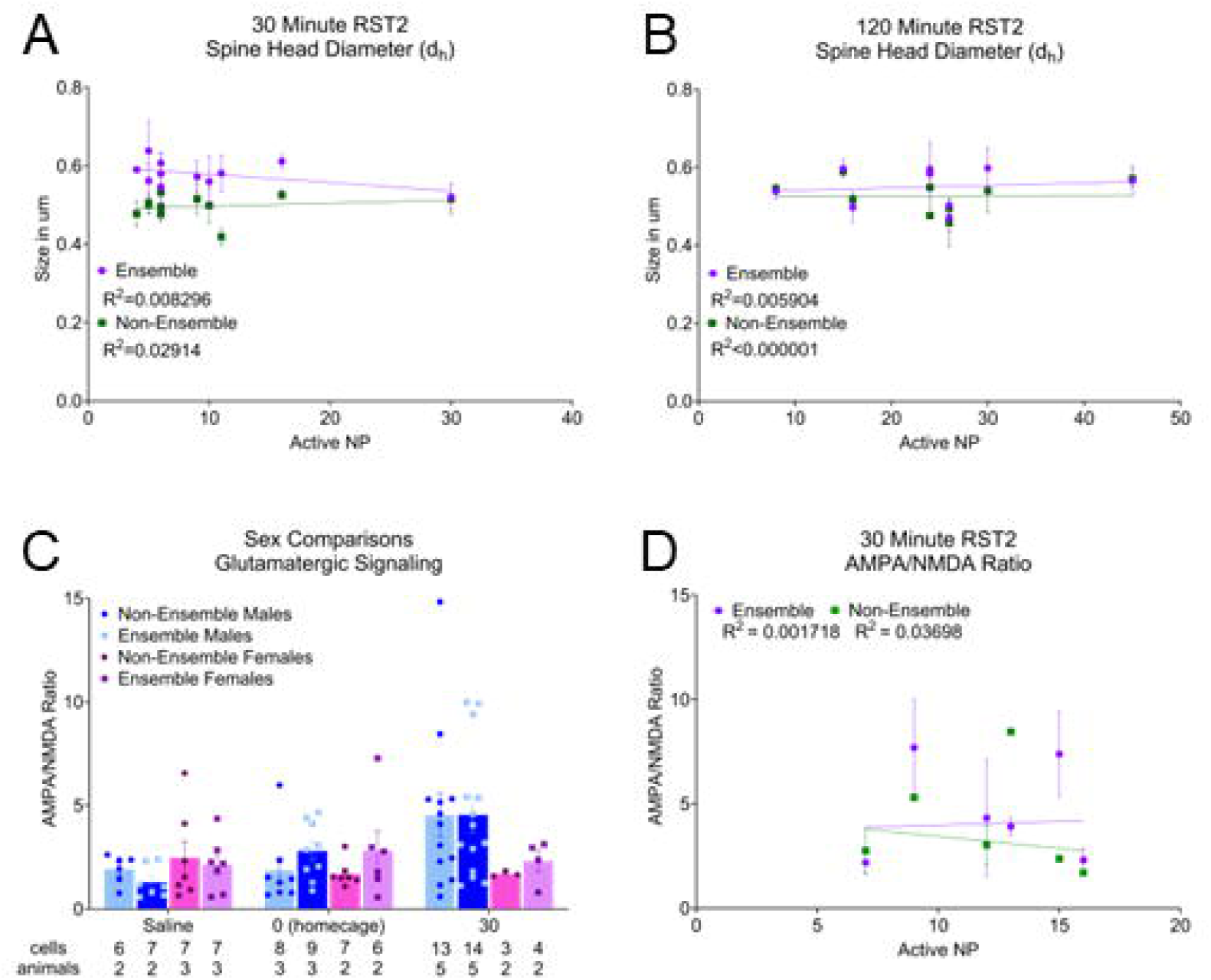
Spine head diameter (dh) and AMPAI NMDA ratio does not correlate with seeking on reinstatement (RST)2 (**A**) Correlation of 30-minute RST2 to dh. (**B**) Correlation of 120-minute RST2 to dh. (**C**) sex comparisons of the AMPAINMDA ratio. (**D**) Correlation of 30-Minute RST2 to cell population AMPAINMDA ratio.

**Extended Data Figure 5.**
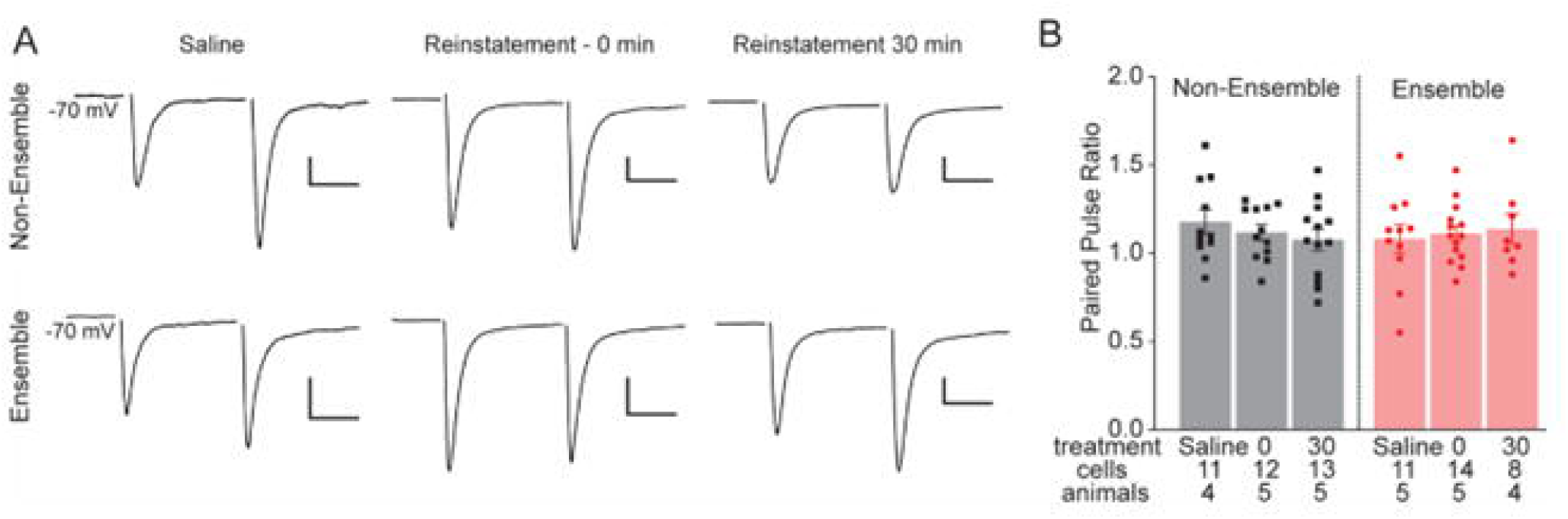
Paired pulse ratio of ensemble and non-ensemble neurons at different timepoints (**A**) Representative traces of paired pulse data from all cell types and time points Scalebars 200 pA/20ms. (**B**) No difference in paired pulse ratio regardless of cell type and time point.

## References

Adinoff B (2004) Neurobiologic Processes in Drug Reward and Addiction. Harvard Review of Psychiatry 12:305–320.

Bobadilla A-C, Dereschewitz E, Vaccaro L, Heinsbroek JA, Scofield MD, Kalivas PW (2020) Cocaine and sucrose rewards recruit different seeking ensembles in the nucleus accumbens core. Mol Psychiatry 25:3150–3163.

Bossert JM, Stern AL, Theberge FRM, Cifani C, Koya E, Hope BT, Shaham Y (2011) Ventral medial prefrontal cortex neuronal ensembles mediate context-induced relapse to heroin. Nat Neurosci 14:420–422.

Brown JA, Petersen N, Centanni SW, Jin AY, Yoon HJ, Cajigas SA, Bedenbaugh MN, Luchsinger JR, Patel S, Calipari ES, Simerly RB, Winder DG (2023) An ensemble recruited by α2a-adrenergic receptors is engaged in a stressor-specific manner in mice. Neuropsychopharmacol 48:1133–1143.

Chan A, Willard A, Mulloy S, Ibrahim N, Sciaccotta A, Schonfeld M, Spencer SM (2022) Metformin in nucleus accumbens core reduces cue-induced cocaine seeking in male and female rats. Addict Biol 27:e13165.

Christian DT, Wang X, Chen EL, Sehgal LK, Ghassemlou MN, Miao JJ, Estepanian D, Araghi CH, Stutzmann GE, Wolf ME (2017) Dynamic Alterations of Rat Nucleus Accumbens Dendritic Spines over 2 Months of Abstinence from Extended-Access Cocaine Self-Administration. Neuropsychopharmacology 42:748–756.

Cruz FC, Rubio FJ, Hope BT (2015) Using c-fos to study neuronal ensembles in corticostriatal circuitry of addiction. Brain Res 1628:157–173.

DeNardo LA, Liu CD, Allen WE, Adams EL, Friedmann D, Fu L, Guenthner CJ, Tessier-Lavigne M, Luo L (2019) Temporal Evolution of Cortical Ensembles Promoting Remote Memory Retrieval. Nat Neurosci 22:460–469.

Garcia-Keller C, Scofield MD, Neuhofer D, Varanasi S, Reeves MT, Hughes B, Anderson E, Richie CT, Mejias-Aponte C, Pickel J, Hope BT, Harvey BK, Cowan CW, Kalivas PW (2020) Relapse-Associated Transient Synaptic Potentiation Requires Integrin-Mediated Activation of Focal Adhesion Kinase and Cofilin in D1-Expressing Neurons. J Neurosci 40:8463–8477.

Gipson CD, Kupchik YM, Kalivas PW (2014) Rapid, Transient Synaptic Plasticity in Addiction. Neuropharmacology 76:10.1016/j.neuropharm.2013.04.032.

Gipson CD, Kupchik YM, Shen H, Reissner KJ, Thomas CA, Kalivas PW (2013a) Relapse induced by cues predicting cocaine depends on rapid, transient synaptic potentiation. Neuron 77:867–872.

Gipson CD, Reissner KJ, Kupchik YM, Smith ACW, Stankeviciute N, Hensley-Simon ME, Kalivas PW (2013b) Reinstatement of nicotine seeking is mediated by glutamatergic plasticity. Proc Natl Acad Sci U S A 110:9124–9129.

Graziane NM, Sun S, Wright WJ, Jang D, Liu Z, Huang YH, Nestler EJ, Wang YT, Schlüter OM, Dong Y (2016) Opposing mechanisms mediate morphine- and cocaine-induced generation of silent synapses. Nat Neurosci 19:915–925.

Greenberg ME, Ziff EB (1984) Stimulation of 3T3 cells induces transcription of the c-fos proto-oncogene. Nature 311:433–438.

Grimm JW, Lu L, Hayashi T, Hope BT, Su T-P, Shaham Y (2003) Time-Dependent Increases in Brain-Derived Neurotrophic Factor Protein Levels within the Mesolimbic Dopamine System after Withdrawal from Cocaine: Implications for Incubation of Cocaine Craving. J Neurosci 23:742–747.

Guenthner CJ, Miyamichi K, Yang HH, Heller HC, Luo L (2013) Permanent Genetic Access to Transiently Active Neurons via TRAP: Targeted Recombination in Active Populations. Neuron 78:773–784.

Hebb DO (1949) The organization of behavior; a neuropsychological theory. Oxford, England: Wiley.

Henton WW, Fisher BR (1981) Cyclical Food Deprivation and Classical Conditioned Response Patterns. Psychol Rec 31:377–393.

Hilderbrand ER, Lasek AW (2014) Sex differences in cocaine conditioned place preference in C57BL/6J mice. Neuroreport 25:105–109.

Horenstein BR (1951) Performance of conditioned responses as a function of strength of hunger drive. Journal of Comparative and Physiological Psychology 44:210–224.

Huang YH, Lin Y, Mu P, Lee BR, Brown TE, Wayman G, Marie H, Liu W, Yan Z, Sorg BA, Schlüter OM, Zukin RS, Dong Y (2009) In vivo Cocaine Experience Generates Silent Synapses. Neuron 63:40–47.

Koob GF, Le Moal M (2001) Drug Addiction, Dysregulation of Reward, and Allostasis. Neuropsychopharmacol 24:97–129.

Koob GF, Volkow ND (2010) Neurocircuitry of Addiction. Neuropsychopharmacology 35:1051.

Koya E, Cruz FC, Ator R, Golden SA, Hoffman AF, Lupica CR, Hope BT (2012) Silent synapses in selectively activated nucleus accumbens neurons following cocaine-sensitization. Nat Neurosci 15:1556–1562.

Koya E, Golden SA, Harvey BK, Guez-Barber DH, Berkow A, Simmons DE, Bossert JM, Nair SG, Uejima JL, Marin MT, Mitchell TB, Farquhar D, Ghosh SC, Mattson BJ, Hope BT (2009) Targeted disruption of cocaine-activated nucleus accumbens neurons prevents context-specific sensitization. Nat Neurosci 12:1069–1073.

Ku MJ, Kim CY, Park JW, Lee S, Jeong EY, Jeong J-W, Kim WY, Kim J-H (2024) Wireless optogenetic stimulation on the prelimbic to the nucleus accumbens core circuit attenuates cocaine-induced behavioral sensitization. Neurobiology of Disease 203:106733.

Lee BR, Ma Y-Y, Huang YH, Wang X, Otaka M, Ishikawa M, Neumann PA, Graziane NM, Brown TE, Suska A, Guo C, Lobo MK, Sesack SR, Wolf ME, Nestler EJ, Shaham Y, Schlüter OM, Dong Y (2013) Maturation of silent synapses in amygdala-accumbens projection contributes to incubation of cocaine craving. Nature Neuroscience 16:1644–1651.

Lenoir M, Engeln M, Navailles S, Girardeau P, Ahmed SH (2024) A large-scale c-Fos brain mapping study on extinction of cocaine-primed reinstatement. Neuropsychopharmacology 49:1459–1467.

Leslie JC (1977) Effects of Food Deprivation and Reinforcement Magnitude on Conditioned Suppression. Journal of the Experimental Analysis of Behavior 28:107–115.

Litif CG, Flom LT, Sandum KL, Hodgins SL, Vaccaro L, Stitzel JA, Blouin NA, Mannino MC, Gigley JP, Schoborg TA, Bobadilla A-C (2023) Sex-Dependent Genetic Expression Signatures within Cocaine- and Sucrose-Seeking Ensembles in Mice. bioRxiv:2023.11.02.565378.

Lynch WJ, Carroll ME (2000) Reinstatement of cocaine self-administration in rats: sex differences. Psychopharmacology 148:196–200.

Mongrédien R et al. (2025) Astrocytes Control Cocaine-Induced Synaptic Plasticity and Reward Through the Matricellular Protein Hevin. Biological Psychiatry 98:612–623.

Roberts-Wolfe D, Bobadilla A-C, Heinsbroek JA, Neuhofer D, Kalivas PW (2018) Drug Refraining and Seeking Potentiate Synapses on Distinct Populations of Accumbens Medium Spiny Neurons. J Neurosci 38:7100–7107.

Roberts-Wolfe DJ, Heinsbroek JA, Spencer SM, Bobadilla AC, Smith ACW, Gipson CD, Kalivas PW (2020) Transient synaptic potentiation in nucleus accumbens shell during refraining from cocaine seeking. Addict Biol 25:e12759.

Roy DS, Park Y-G, Kim ME, Zhang Y, Ogawa SK, DiNapoli N, Gu X, Cho JH, Choi H, Kamentsky L, Martin J, Mosto O, Aida T, Chung K, Tonegawa S (2022) Brain-wide mapping reveals that engrams for a single memory are distributed across multiple brain regions. Nat Commun 13:1–16.

Sebastian V, Estil JB, Chen D, Schrott LM, Serrano PA (2013) Acute Physiological Stress Promotes Clustering of Synaptic Markers and Alters Spine Morphology in the Hippocampus. PLOS ONE 8:e79077.

Shen H, Moussawi K, Zhou W, Toda S, Kalivas PW (2011) Heroin relapse requires long-term potentiation-like plasticity mediated by NMDA2b-containing receptors. Proc Natl Acad Sci U S A 108:19407–19412.

Sortman BW, Gobin C, Rakela S, Cerci B, Warren BL (2022) Prelimbic Ensembles Mediate Cocaine Seeking After Behavioral Acquisition and Once Rats Are Well-Trained. Frontiers in Behavioral Neuroscience 16 Available at: https://www.ncbi.nlm.nih.gov/pmc/articles/PMC9549214/ [Accessed January 3, 2023].

Wang W, Xie X, Zhuang X, Huang Y, Tan T, Gangal H, Huang Z, Purvines W, Wang X, Stefanov A, Chen R, Rodriggs L, Chaiprasert A, Yu E, Vierkant V, Hook M, Huang Y, Darcq E, Wang J (2023) Striatal μ-opioid receptor activation triggers direct-pathway GABAergic plasticity and induces negative affect. Cell Reports 42:112089.

Wang YQ, Huang YH, Balakrishnan S, Liu L, Wang YT, Nestler EJ, Schlüter OM, Dong Y (2021) AMPA and NMDA Receptor Trafficking at Cocaine-Generated Synapses. J Neurosci 41:1996–2011.

Whitaker LR, Carneiro de Oliveira PE, McPherson KB, Fallon RV, Planeta CS, Bonci A, Hope BT (2016) Associative learning drives the formation of silent synapses in neuronal ensembles of the nucleus accumbens. Biol Psychiatry 80:246–256.

Whitaker LR, Hope BT (2018) Chasing the addicted engram: identifying functional alterations in Fos-expressing neuronal ensembles that mediate drug-related learned behavior. Learn Mem 25:455–460.

Williamson MR, Kwon W, Woo J, Ko Y, Maleki E, Yu K, Murali S, Sardar D, Deneen B (2025) Learning-associated astrocyte ensembles regulate memory recall. Nature 637:478–486.

Wolf ME (2016) Synaptic mechanisms underlying persistent cocaine craving. Nat Rev Neurosci 17:351–365.

Wright WJ, Dong Y (2021) Silent Synapses in Cocaine-Associated Memory and Beyond. J Neurosci 41:9275–9285.

Wright WJ, Graziane NM, Neumann PA, Hamilton PJ, Cates HM, Fuerst L, Spenceley A, MacKinnon-Booth N, Iyer K, Huang YH, Shaham Y, Schlüter OM, Nestler EJ, Dong Y (2020) Silent Synapses Dictate Cocaine Memory Destabilization and Reconsolidation. Nat Neurosci 23:32–46.

